# Diversity and distribution of the coral-associated endolithic algae *Ostreobium* in the Southwestern Caribbean

**DOI:** 10.1101/2023.05.18.541270

**Authors:** A.P. Rodríguez-Bermúdez, S. Ramírez-Palma, J.S. Giraldo-Vaca, L.M. Diaz-Puerto, J.A Sánchez

## Abstract

Coral reefs are facing significant environmental challenges. Ocean acidification has the potential to induce the dissolution of coral reefs. The community of micro-bioerosion exhibits a heightened level of concern in the context of ocean acidification. Comprehending the close interplay between bioeroders and corals is of utmost importance in predicting the trajectory of these vulnerable ecosystems. The genus *Ostreobium*, which belongs to the order Bryopsidales (Chlorophyta) and comprises euendolithic chlorophyte algae, has been identified as the primary cause of reef dissolution among microbioeroders. The objective of this study was to comprehend the inherent distribution of *rbcL* clades of *Ostreobium* in the Southwestern Caribbean corals within a gradient of reef depth. The *Ostreobium rbcL* variants were found to be linked with corals belonging to the Agariciidae, Merulinidae, Poritiidae, Siderastreidae, Astroconeiidae, Montastreidae, Mussidae, and Pocilloporidae families. The individuals selected for the current investigation are categorized as members of either the “complex” or “robust” coral groups. Our findings indicate that solely the corals categorized as ‘complex’ exhibit close memberships with the three *Ostreobium* superclades. In general, the dispersion of *Ostreobium* within the Southwestern Caribbean region exhibits differentiation among various coral groups and is influenced by geographical and bathymetric factors. The *Ostreobium*’s diversity is primarily composed of ecological specialists, wherein most clades are linked to particular hosts. Conversely, only a few ecological generalists are associated with multiple hosts, akin to zooxanthellae. *Ostreobium* exhibits greater diversity on encrusting corals such as agariciids, which are among the most abundant and widespread coral species in the Caribbean.

## INTRODUCTION

Coral reefs are under unprecedented environmental threats. Ocean thermal anomalies cause coral bleaching and mortality (Hughes et al. 2003; Eakin et al. 2010) and ocean acidification has a negative impact on the calcification rates of marine organisms such as corals (Orr et al. 2005; Hoegh-Guldberg et al. 2007). This process interacts with the human footprint, specifically growing sewage and overexploitation, reducing the resilience of marine ecosystems like coral reefs (Graham et al. 2008, 2013; Wiedenmann et al. 2013). Particularly, ocean acidification may cause reef dissolution (Hoegh-Guldberg et al. 2017). Scleractinian corals, the main reef-building organisms, had already shown a decrease in calcification rates (De’ath et al. 2009) in response to a reduction in Ω_arag_ saturation (Fantazzini et al. 2015). Of particular concern, the micro-bioerosion community increases under conditions of ocean acidification (Enochs et al. 2015). Understanding the tight interaction between bioeroders and corals is critical for forecasting the future of these fragile ecosystems.

The euendolithic chlorophyte algae genus *Ostreobium*, a siphonous green algae from the order Bryopsidales (Chlorophyta), is the most prevalent agent responsible for reef dissolution among microbioeroders (Tribollet 2008; Tribollet et al. 2009; Grange et al. 2015). Ostreobium has also gained popularity due to its ability to undertake low-light photosynthesis, which allows it to spread further into the depths (Rouzé et al., 2021; Verbruggen & Tribollet, 2011). Furthermore, the presence of chlorophyll b and lutein in *Ostreobium* has perplexed scientists given the absence of these pigments in the most common group of endosymbiotic dinoflagellate Symbiodinaceae (Jeffrey 1968a, 1968b; Apprill et al. 2007). The adaptive ecophysiology of *Ostreobium* is related to the rocky and harsh environment in which it dwells (Ricci et al. 2019), its interaction with the coral holobiont during bleaching events (Fine et al. 2004, 2006a; Galindo-Martínez et al. 2022) and a closed connection to coral tissues at mesophotic depths (Gonzalez-Zapata et al. 2018b). Whereas those research opportunities and challenges have been addressed to some extent, the distribution of individual genotypes in natural habitats remains poorly understood (Tandon et al., 2023). It plays a dynamic role within the skeleton of scleractinian corals (Bornet and Flahault 1889a), but limited information exists on aspects as important as their substrate preference or role as symbiont (Kobluk and Risk 1977; Grange et al. 2015). During bleaching, heat stress, and disease events, *Ostreobium* utilises coral metabolic waste to translocate fixed carbon to the coral host, indicating a clear transition from a commensalistic to a mutualistic relationship (Fine and Loya 2002; Fine 2005; Fine et al. 2006b). A mutualistic relationship with bioeroders such as *Ostreobium* may allow corals to coexist with bioerosion.

A defining trait of reef-building coral species is the ensemble of symbiotic linkages with a microbial community, which plays a crucial role in determining the robustness of its coral host and, by extension, the ecosystem (Bourne et al. 2016; Peixoto et al. 2017). While the symbiosis between corals and zooxanthellae is the most well-studied within the coral holobiont, the dynamics of other microorganisms, such as those inhabiting the coral skeleton, have received little attention (Försterra and Häussermann 2008). Taxonomically, *Ostreobium* has three valid species: *O. queketii* (Bornet and Flahault 1889b), 1889, *O constrictum* (Lukas 1974), *O. reineckii* (Bornet and Flahault 1889b). However, only a few morphological characters are diagnostic, limiting morphological approaches and making species differentiation difficult. A recent study that used the *rbcL* barcode identifier discovered more species than previous taxonomic literature indicated (Gutner-Hoch and Fine 2011). Since then, the molecular data obtained through the sequencing of plastid encoded markers such as *rbcL, tufA*, UPA, and 16S rRNA increased and allowed us understand better the prevalence of *Ostreobium* in the core microbiome of tropical corals, along with its extensive genetic diversity (Gutner-Hoch and Fine 2011; Marcelino and Verbruggen 2016; del Campo et al. 2017; Gonzalez-Zapata et al. 2018b). Until now, we are grasping on to the natural distribution of *Ostreobium* diversity. It is potentially associated in nearly 85% of coral species (Gutner-Hoch and Fine 2011), on a wide geographical range (del Campo et al. 2017) and bathymetric range (deepest record at 172 m, (Rouzé et al. 2021). Here, we seek to understand the natural distribution of *rbcL* clades of *Ostreobium* in the Southwestern Caribbean in a reef depth gradient.

## METHODS

### Field sampling

We collect 200 coral samples at two localities in the Colombian Caribbean: Cartagena de Indias and San Andres Island between December 2019 and March 2020, and during the months of March 2021 and 2022 (Fig. 1). Subsamples of 1cm^2^ were preserved in ETOH 96% and stored at −80°C until DNA extraction. The coral colonies were identified using the taxonomic keys of (Veron 2000; Wells). A dry voucher is available at the Museo de Historia Natural Uniandes (Supplementary Table 1). Research and collection of specimens were approved by the National Environmental Licensing Authority (ANLA, Spanish acronym): Collection Framework Agreement granted to Universidad de los Andes through resolutions No. 02215 of 8^th^ November 2019, and agreements with Parques Nacionales and Oceanario-CEINER Islas del Rosario (Observatorio de Microbioerosion Marina), 2022.

**Figure 1.**
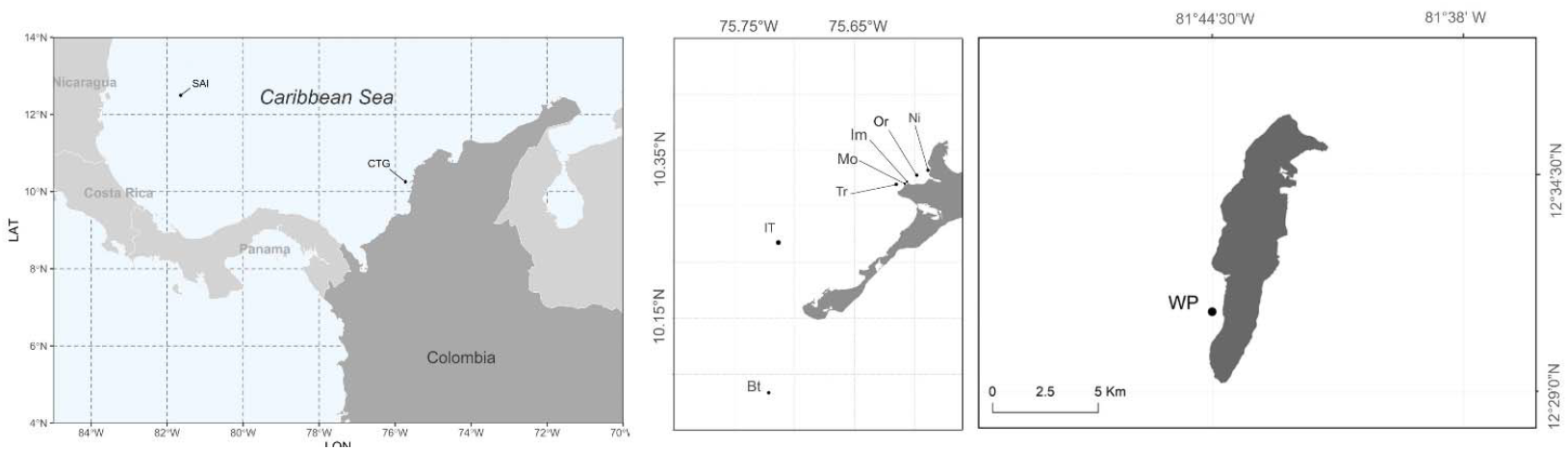
Sampling localities. Study area in the Colombian Caribbean. Stations at Cartagena de Indias (CTG): Ni = Nico, OR = Octubre Rojo, Tr = Trompadas, Mo = Montañita, Im = Imelda, IT = Isla Tesoro, Bt = Bajo Tortugas; San Andres Island (SAI): WP = Western Point.

### Molecular detection of *Ostreobium*

To prepare the sample for DNA extraction, we washed a small part of the coral colony (∼0.5cm^2^) preserved in ETOH 96% with a WaterPick© using a pressurized jet of MiliQ water to remove the coral tissue. Then, we exposed the green band in the skeleton belonging to the Ostreobium algae community with a MotorTool and powder grounded using a mortar and pestle. Finally, we proceed to extract DNA following the CTAB protocol used in (Cremen et al. 2016).

### *rbcL* Amplification

We amplified a ∼430bp fragment of the chloroplast encoded *Ostreobium* rbcL gene using the primers: rcbL250 [5’GATATTGARCCTGTTGTTG GTGAAGA 3 ‘] and rcbLR670 [5’ CCAGTTTCAGCTTGWGCTTTATAAA 3’] (Massé et al. 2020). We adjust the PCR reactives to following: 15ul reaction containing 3ul[5X] OneTaq reaction buffer, 0.6ul of 10mM forward and reverse primes, 0.3ul [1 U] One Taq DNA polymerase (Biolabs, USA), 0.3 [10mM] DNTPs, 2.1 [1.5mM] MgCl++, 0.75ul[20mg/ul] BSA, 6.35 Milli Q water and 1ul[30ng] template DNA. The PCR conditions were denaturation at 94°C for 2min, annealing for 35 cycles at 94°C for 90s, 55°C for 90s and 72°C for 60s-, and 5-min extension at 72°C. We visualized the amplified fragments in 1.5% agarose gel with SYBR™Gold. PCR products were cleaned with FastAP Thermosensitive Alkaline Phosphatase and sequenced using the BigDye Terminator v3.1 Cycle Sequencing Kit (Applied Biosystems) on the AD1373xl DNA Analyzer (Applied Biosystems). Finally, we sent the positive PCR products to GenCore lab to do Sanger sequencing. Out of 200 samples, a total of 115 samples amplified for the *rbcL* gene, 63 sequences had enough quality to perform further phylogenetic analysis (31% of the samples). Both ML and Bayesian consensus trees conserved the same topologies, only Bayesian inferred tree was shown (Figure 2.A).

**Figure 2.**
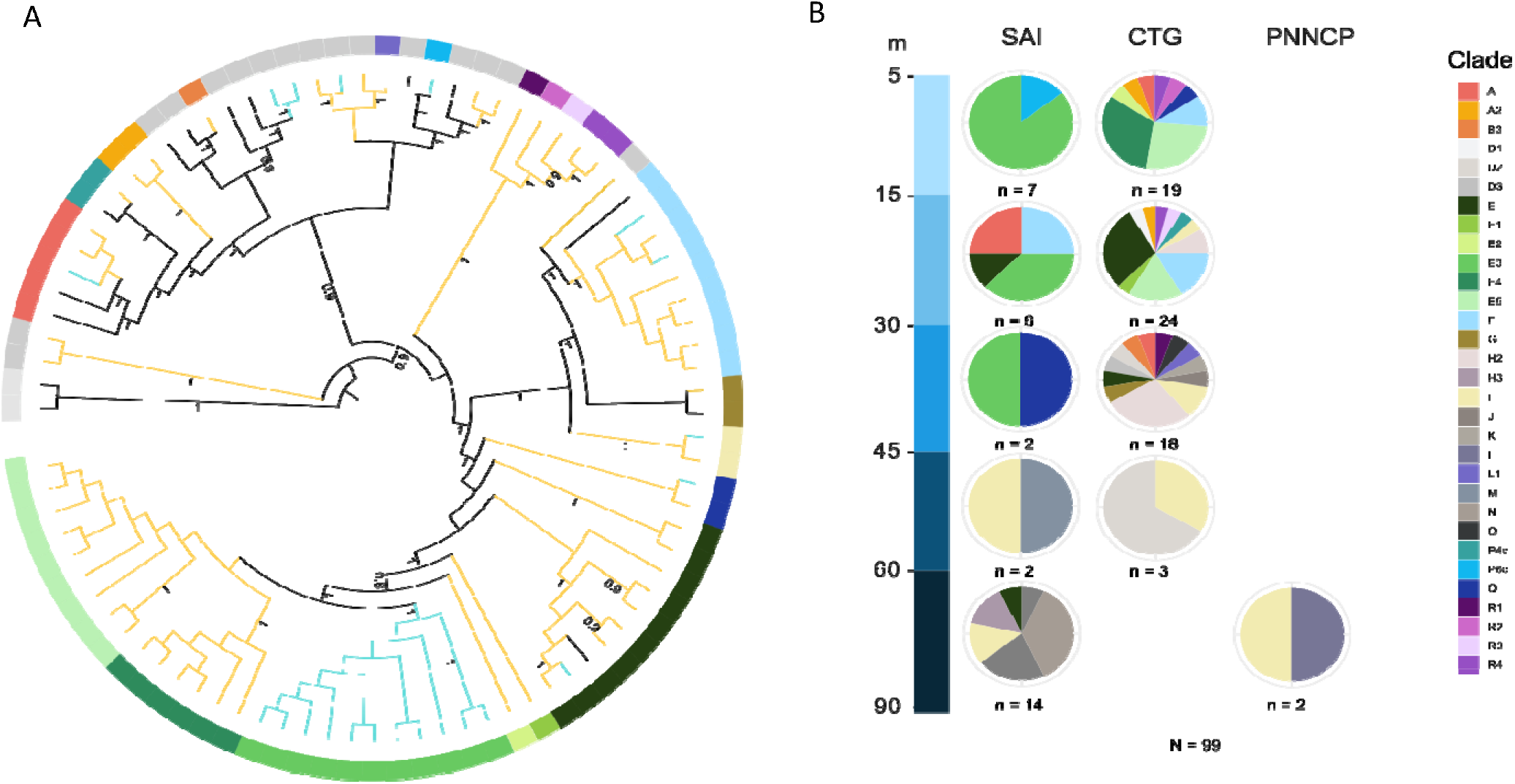
**A**. Summarized in *rbcL* clades bayesian phylogenetic tree of *Ostreobium*. Bootstrap values > 0.8 are shown. Branch colors correspond to localities in the Caribbean: cyan color (SAI), orange yellow (CTG), and black to other locations outside the Caribbean. The circle band outside the tree indicates the species delimitation, light-gray color at the band correspond to *rbcL* clades identified in previous studies. (See details in Supplementary Table 2). **B**. Depth distribution of *Ostreobium rbcL* clades reported for the Colombian Caribbean. We included data from the National Natural Park Deep-Sea Corals (PNNCP) to analyze the genetic diversity outlook of *Ostreobium* in the Caribbean using metadata from Gonzalez-Zapata et al., (2018).

### Phylogenetic analysis

We aligned new sequences accession numbers from OQ935479 to OQ935534 (Supplementary Table 1), NCBI *Ostreobium rbcL* sequences from previous studies (Supplemetary Table 2), and two outgroups from the Bryopsidales order used by (Iha et al. 2021). The multiple sequence alignment was performed using the ClustalW algorithm in MEGA11 (Kumar et al. 2018). We estimated maximum likelihood trees in RAXMLv8(Stamatakis, 2014) using the GTR model and 1000 bootstrap replicates. We also estimated ultrametric Bayesian trees using BEAST v2.5. We used JModelTest2 (Darriba et al. 2012) to estimate the best fitting substitution model according to the corrected Akaike information criterion (Lukas 1974; Sugiura 1978). In BEAST, we applied the resulting TrN+I+G model, an Optimised Relaxed Clock, Birth Death Model of speciation prior, and ran analyses for 10 million Markov chain Monte Carlo (MCMC) generations. We confirmed the resulting.log files in TRACER v1.4 (Drummond and Rambaut 2007), and found that each analysis reached stationarity and had effective sample size (ESS) values >200. Finally, we discarded the first 5000 trees (10%) as burn-in. Resulting trees were annotated in TreeAnnotator.

### Clades delimitation

We implemented the two following methods to identify *Ostreobium* clades: a generalized mixed Yule coalescent model using the function gmyc in the R package Splits (Fujisawa and Barraclough 2013) and the coalescent Poisson tree process (bPTP) with Bayesian support (BS) method (Zhang et al. 2013) with the parameters 100.000 MCMC generations and a 0.1 burn-in factor. To avoid the hyperinflation and complex interpretation of PTP and GMYC methods in delimiting single-gene trees, we interpreted the resulting clades alongside ecological features such as geography and depth, and named according previous studies(Gutner-Hoch and Fine 2011; Gonzalez-Zapata et al. 2018b; Massé et al. 2020). Consensus trees were visualized and modified in FigTree v1.4.4. and iTOL website (http://itol.embl.de/).

## RESULTS

### Phylogeny of the Southwestern Caribbean Ostreobium

Phylogenetic analyses of the chloroplast *rbcL* region (∼ 410 bp) revealed a community of 20 well-supported clades. 15 new clades reported in the present study (clades A2, B3, E1, E2, E3, E4, E5, L1, P4c, P6c, Q, R1, R2, R3, R4) and three new reports for the Colombian Caribbean region (clades A, F and G) (Figure 2.A).

### Geographical structure of Ostreobium

The principal clades of *Ostreobium* were structured by geography. Clade OstA comprises worldwide clades, while clade OstB includes clades from the Red-Sea and the Caribbean. Finally, clade OstC is almost exclusively composed of Colombian Caribbean clades, except for one genotype from the Red Sea (See also Supplementary Figure A). Within clade OstC, we found geographical structure by locality (SAI or CTG). In terms of location specificity and dominance, only the clades A, E, F, I and Q were shared between CTG and SAI. The clades E3, M and N are abundant and only found at SAI. At CTG, 14 clades are exclusive of this location, increasing the reported diversity of *Ostreobium* to 20 different clades, making it the most diverse locality.

### Depth-structured patterns

Our results showed that all three ‘superclades’ OstA, OstB and OstC are represented at all the depth ranges of the analysis (present study and supplementary data from Gonzalez-Zapata et al. 2018). However, figure 2.B. shows some depth-related distribution patterns: the ‘depth-generalists’ and ‘depth-specific clades’. Among the depth-generalists, we highlight the presence of clades E, E2, H2 and I, which are present from shallower waters (<30m) to mesophotic zones (30m −90m). We also found clades A, A2, F, R2 and Q present across all the ranges but limited until the upper mesophotic zone (<45m). While clades L1, E2, E4 and E5 are restricted to shallow waters (<30m), clades L, M, N and H3 remain restricted to deeper zones (>30m), as previously reported (FG2018). At CTG the depth range with higher diversity of *Ostreobium* is the 30 to 45m depth with thirteen different clades, while in SAI, the depth most diverse range is deeper (60 – 90 m) with five clades.

### Ostreobium hosts

We successfully amplified *rbcL Ostreobium* types associated with corals from the families Agariciidae, Merulinidae, Poritiidae, Siderastreidae, Astroconeiidae, Montastreidae, Mussidae and Pocilloporidae. The representatives of each in the present study belong to either ‘complex’ or ‘robust’ coral groups. We found that only ‘complex’ corals have associations with all three *Ostreobium* superclades, and the ‘robust’ corals mostly associate with OstC (Figure 3.A). We also found a high diversity associated to Agariciids (24 clades out of 31, 77% of known *Ostreobium* clades for the Colombian Caribbean), followed by Poritiids (7 out 31, 22%) and finally Meruliniids (4 out 31, 12%). While Agariciids are associated with the three superclades of *Ostreobium*, Poritiids are associated only with OstB and OstA, and Meruliniids are associatedexclusively with OstC. So far, the clades I, E3 E4, E5, R4, H2, H3, M, N, O, and D are reported only in association with Agariciids despite locality and depths. Two new clades, namely P4c and P6c, have been identified exclusively in *Porites colonensis*. These clades are sister taxa of cultured strains isolated from Indo-Pacific Pocilloporids (See P4 and P6 clades at Massé et al. 2018). Their position at the phylogenetic tree matches the findings reported in (Massé et al. 2018), and also, are found in *Porites* colonies suggesting a possible relation between hosts and associated type of *Ostreobium*.

**Figure 3.**
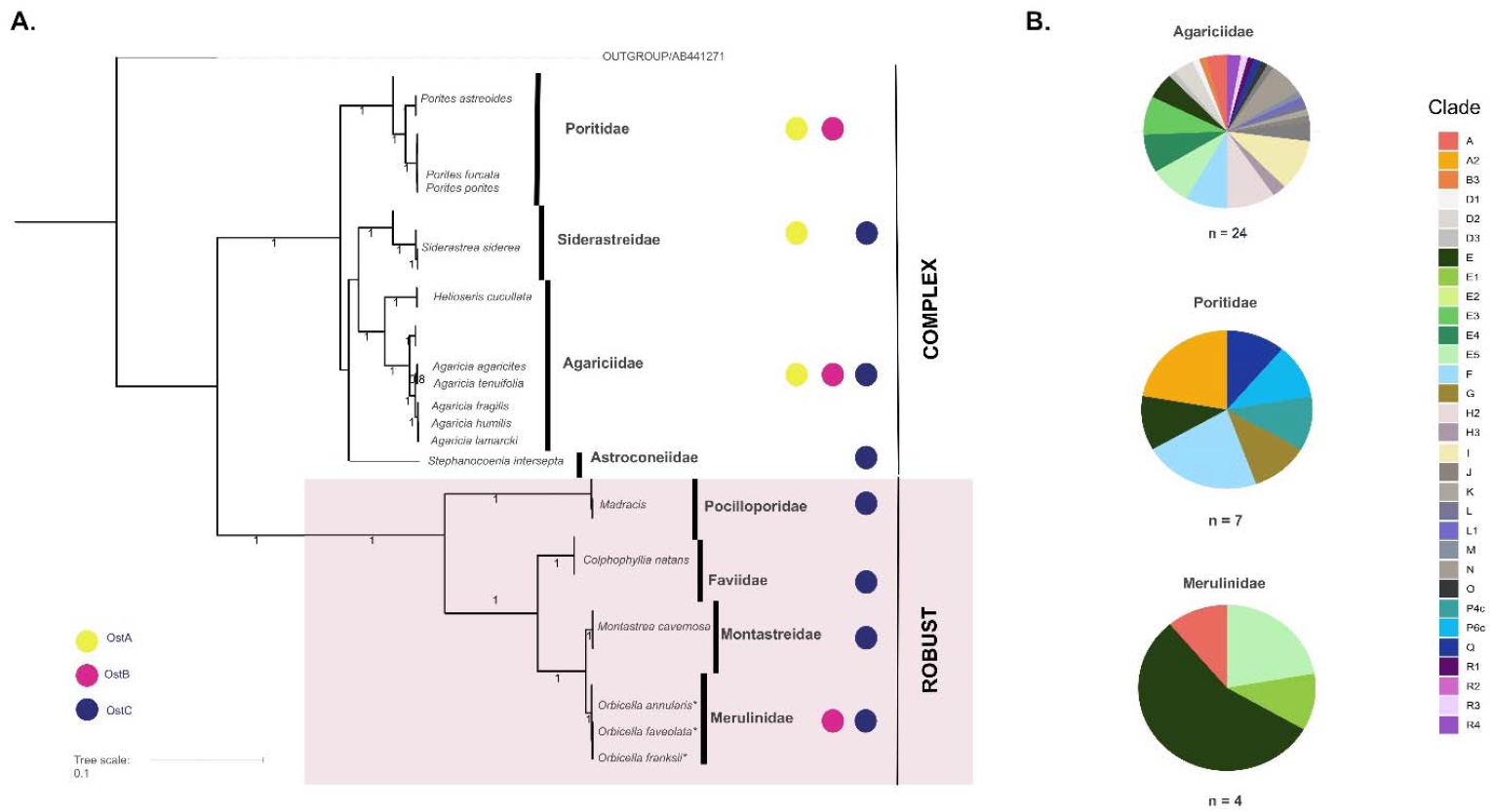
**A**. Distribution of *Ostreobium* principal clades in the coral families assessed in the Colombian Caribbean based on *rbcL* sequences. The coral phylogenetic tree reconstructed from selected data of the CO1 marker from (Kitahara et al. 2010). Only the coral species included in the present study were annotated in the tree labels, species without CO1 information were assumed to share the same phylogenetic position with species on the same genus. The three *Ostreobium* superclades where labeled at the left corner. B. Scleractinian families with the highest number of *Ostreobium* individual clades.

## DISCUSSION

Our study reveals that the spatial distribution of *Ostreobium* in the Southwestern Caribbean exhibits differentiation across various coral groups and is subject to the influence of geographical and bathymetric factors. The concept of environmental heterogeneity in a coral-endosymbiont relationship, which encompasses variations in both biotic and abiotic factors across space and time, is widely recognized as a fundamental determinant of species richness patterns at various scales, spanning from local to continental levels (LaJeunesse and Thornhill 2011; Grajales and Sanchez 2016). *Ostreobium* diversity depends on the location and symbiont availability (del Campo et al., 2017). The study presented additional empirical support indicating that various rbcL clades (A, F, G, I, and E) exhibit a wider distributional range due to the vast expanse of the Caribbean and Red Sea, which are major marine basins situated in the Atlantic and Indian Oceans, respectively. Moreover, the aforementioned clades exhibit distinct depth structures and varying relationships with numerous hosts.

### Geographic and depth zonation

Del Campo et al. (2017) indicated the presence of the three dominant *Ostreobium* groups defined by the 16S rRNA in the Caribbean. Similarly, we found three major clades OstA, OstB, OstC with similar distributions (Supplementary Figure 1). It could imply that the *rbcL* marker is a proxy for diversity and comparable to 16s RNA-generated topologies. So far, the *rbcL* marker has revealed the highest reported diversity for *Ostreobium* in the Caribbean region. Differences in clade composition between SAI and CTG locations can be attributed to the environmental heterogeneity among sites such as reef setting (oceanic: SAI vs continental influence: CTG), light regimens, turbidity and predators (Velásquez and Sánchez 2015), all of which have a direct impact on the physiology and population persistence of *Ostreobium*.

The ‘insurance theory’ could explain the distribution pattern of *Ostreobium* in Cartagena in particular. According to this idea, ecosystem stability is increased by diversity and functional redundancy due to complementarity and asynchronous reactions to stress (Loreau and de Mazancourt 2013). This idea is similar to the adaptive dysbiosis hypothesis in that microorganisms give resistance and improve host resilience due to functional redundancy (Boilard et al. 2020). Because *Ostreobium* is better adapted to extreme environments, it is not surprising to find higher clade diversity under the dim light conditions that characterize Cartagena reefs (Roitman et al. 2020), where continental run-off reduces the penetration of light (Gonzalez-Zapata et al. 2018a). Our findings in this regard contrast those of Symbiodiniaceae, where diversity depletion occurs on turbid zones (López-Londoño et al. 2021). These conditions are suitable for green algae like *Ostreobium*, increasing its diversity (del Campo et al. 2017). Presumably, some better adapted *Ostreobium* clades may replace some zooxanthellae functions in environments with stressful conditions, easing coral acclimation to bleaching stress and light scarcity. However, it is unclear how this diversity aids in the holobiont’s adaptation to those specific conditions.

### The influence of Host species Identity

In contrast to what del Campo et al. (2017) found at the Caribbean, our findings show that ‘robust’ corals are associated with fewer types of *Ostreobium* than ‘complex’ corals. While this estimation can be hindered by the large number of Agariciids samples, it can also tell us how the dominance of *rbcL* clades in the *Ostreobium* community of a particular sample can hide the cryptic biodiversity that could be discovered with metabarcoding approaches that include other markers (such as 16S rRNA or *tufA*).

Similar to zooxanthellae, *Ostreobium* diversity appears to be dominated by “ecological specialists” with the majority of clades associated with specific hosts, whereas few “ecological generalists” are associated with multiple hosts (Finney et al. 2010). *Ostreobium* is more diverse on encrusting corals such as agariciids, one of the most prevalent and prolific coral species in the Caribbean (Gonzalez-Zapata et al. 2018a). Also, agariciids exhibit greater resilience in disturbed habitats like Cartagena, exhibiting high rates of survival after bleaching events (Cáceres and Sánchez 2015; Navas-Camacho et al. 2015), and also is the coral form that inhabits by excellence the reef slope habitats characterized by low-irradiance regimens (Bongaerts et al. 2013; Prata et al. 2022). The diverse repertory of dominant *Ostreobium* clades found at Agariciids illustrates the importance of *Ostreobium* genetic pools in increasing their prevalence and resistance across wide geographic areas. Host species possibly have control on the composition of endolithic community via skeletal specialization (Marcelino et al. 2017). It is remarkable how different the diversity composition is in the most abundant hosts: while clade E is the most prevalent genotype associated with ‘robust’ corals at Cartagena (*Orbicella* spp. and Agariciids species). *Porites* species (‘complex’ corals) exhibited preferences for different and uncommon *Ostreobium* genotypes.

The present study has presented a brief overview of the ecological expansion and diversification of *Ostreobium*. However, further investigation is warranted to explore the *Ostreobium* community profiles associated with a broader range of host species across varying optical characteristics of the water column. Despite recent investigations into the physiological responses of *Ostreobium* in diverse skeletal environments, the adaptive function of *Ostreobium* remains nascent in our comprehension. It is suggested that a proposed framework be employed to enhance the inherent adaptive capacity of coral reefs in response to climate change by investigating the function of supplementary coral holobiont entities, such as *Ostreobium*, as potential targets for adaptive intervention (Voolstra et al. 2021).

## ACKNOWLEDGEMENTS

We thank the BIOMMAR crew for the fruitful discussions during all the stages of this project. To Cartagena divers, Makarela and Sea Pride diving operators that supported us in the field. We especially thank Laura Rodriguez, Sandra Montaño, Andres Chilma, Adrian Devia, Adriana Sarmiento, and Carlos Gomez for being incredible dive buddies and great companionship during the field trip, to Sebastian Guzman for kindly sharing his evolutionary and bioinformatics knowledge through the data interpretation. Finally, to the Universidad de Los Andes, MinCiencias (Ministry of Science, Technology and Innovation), Colombia, for financing the study, within the program “Observatory of microbioerosion, ocean acidification and dissolution in coral reefs” code 1204-852-70251 and Facultad de Ciencias-Uniandes (Proyecto Semilla) for financing the study. We also acknowledge the participation of local communities during the field surveys.

## AUTHORS’ CONTRIBUTIONS

ARB conceived and designed the analysis, collected the data, performed the phylogenetic analysis, conducted part of the field trips and performed laboratory procedures and wrote the manuscript.

JSGV contributed with data, literature review and performed laboratory procedures.

LMD reviewing, editing and writing

SRP did field planning and sampling, DNA extraction and quantification, PCR amplification and reviewed the manuscript.

JAS conceived the study, contributed with data, and reviewed substantially the manuscript.

All authors reviewed the final manuscript.

