## Supplementary material for "Diversity and distribution of the coral-associated endolithic algae *Ostreobium* in the Southwestern Caribbean": Suplementary Material

**Supplementary Table 1**. Metadata of the samples used in the present study.

| **ACCESION NUMBER NCBI** | **VOUCHER** | **FIELD CODE** | **SUPERKINGDOM:KINGDOM:PHYLUM:CLASS:ORDER:FAMILY:GENUS** | **RCBL TYPE** | **HOST PHYLUM:ORDER:FAMILY:GENUS** | **HOST SPECIES** | **COLLECTION DATE** | **LOCALITY** | **LAT_LONG** | **DEPTH** |
| --- | --- | --- | --- | --- | --- | --- | --- | --- | --- | --- |
| OQ935479 | ANDES-IM8201 | BT2 CTG39 | Eukaryota:Viridiplantae:Chlorophyta: Ulvophyceae: Bryopsidales: Ostreobiaceae: Ostreobium | Ostreobium sp.E5 AR-2023 | Cnidaria:Scleractinea:Agariciidae | *Agaricia tenuifolia* | 3/22/2022 | Colombia:Cartagena | 10.09N 75.87W | 15 |
| OQ935480 | ANDES-IM6491 | NIS67 | Eukaryota:Viridiplantae:Chlorophyta: Ulvophyceae: Bryopsidales: Ostreobiaceae: Ostreobium | Ostreobium sp.Q AR-2023 | Cnidaria:Scleractinea:Agariciidae | *Agaricia lamarcki* | 10/9/2017 | Colombia:Cartagena | 10.28N 75.6W | 11 |
| OQ935481 | ANDES-IM8207 | IT CTG51 | Eukaryota:Viridiplantae:Chlorophyta: Ulvophyceae: Bryopsidales: Ostreobiaceae: Ostreobium | Ostreobium sp.E4 AR-2023 | Cnidaria:Scleractinea:Agariciidae | *Agaricia lamarcki* | 3/22/2022 | Colombia:Cartagena | 10.14N 75.44W | 10 |
| OQ935482 | ANDES-IM7884 | CTG21 | Eukaryota:Viridiplantae:Chlorophyta: Ulvophyceae: Bryopsidales: Ostreobiaceae: Ostreobium | Ostreobium sp.R3 AR-2023 | Cnidaria:Scleractinea:Agariciidae | *Helioseris cucullata* | 3/15/2020 | Colombia:Cartagena | 10.26N 75.61W | 15 |
| OQ935483 | ANDES-IM7751 | WP15 | Eukaryota:Viridiplantae:Chlorophyta: Ulvophyceae: Bryopsidales: Ostreobiaceae: Ostreobium | Ostreobium sp.E3 AR-2023 | Cnidaria:Scleractinea:Agariciidae | *Agaricia grahamae* | 5/12/2019 | Colombia:San Andres island | 12.51N 81.73W | 15 |
| OQ935484 | ANDES-IM8199 | BT CTG27 | Eukaryota:Viridiplantae:Chlorophyta: Ulvophyceae: Bryopsidales: Ostreobiaceae: Ostreobium | Ostreobium sp.E5 AR-2023 | Cnidaria:Scleractinea:Agariciidae | *Agaricia fragilis* | 3/22/2022 | Colombia:Cartagena | 10.09N 75.87W | 10 |
| OQ935485 | ANDES-IM8198 | BT CTG26 | Eukaryota:Viridiplantae:Chlorophyta: Ulvophyceae: Bryopsidales: Ostreobiaceae: Ostreobium | Ostreobium sp.E4 AR-2023 | Cnidaria:Scleractinea:Agariciidae | *Agaricia fragilis* | 3/22/2022 | Colombia:Cartagena | 10.09N 75.87W | 10 |
| OQ935486 | ANDES-IM7845 | CTG33 | Eukaryota:Viridiplantae:Chlorophyta: Ulvophyceae: Bryopsidales: Ostreobiaceae: Ostreobium | Ostreobium sp.E AR-2023 | Cnidaria:Scleractinea:Agariciidae | *Agaricia lamarcki* | 3/15/2020 | Colombia:Cartagena | 10.26N 75.62W | 33 |
| OQ935487 | ANDES-IM7492 | I24 | Eukaryota:Viridiplantae:Chlorophyta: Ulvophyceae: Bryopsidales: Ostreobiaceae: Ostreobium | Ostreobium sp.E AR-2023 | Cnidaria:Scleractinea:Merulinidae | *Orbicella faveolata* | 6/14/2018 | Colombia:Cartagena | 10.26N 75.62W | 20 |
| OQ935488 | ANDES-IM7761 | WP26 | Eukaryota:Viridiplantae:Chlorophyta: Ulvophyceae: Bryopsidales: Ostreobiaceae: Ostreobium | Ostreobium sp.E3 AR-2023 | Cnidaria:Scleractinea:Agariciidae | *Helioseris cucullata* | 5/12/2019 | Colombia:San Andres island | 12.51N 81.73W | 30 |
| OQ935489 | ANDES-IM7862 | CTG41 | Eukaryota:Viridiplantae:Chlorophyta: Ulvophyceae: Bryopsidales: Ostreobiaceae: Ostreobium | Ostreobium sp.R1 AR-2023 | Cnidaria:Scleractinea:Agariciidae | *Agaricia undata* | 3/15/2020 | Colombia:Cartagena | 10.26N 75.61W | 33 |
| OQ935490 | ANDES-IM8212 | SH 03 | Eukaryota:Viridiplantae:Chlorophyta: Ulvophyceae: Bryopsidales: Ostreobiaceae: Ostreobium | Ostreobium sp.E3 AR-2023 | Cnidaria:Scleractinea:Mussidae | *Colpophyllia natans* | 3/22/2022 | Colombia:San Andres island | 12.51N 81.73W | 7 |
| OQ935491 | ANDES-IM7870 | CTG17 | Eukaryota:Viridiplantae:Chlorophyta: Ulvophyceae: Bryopsidales: Ostreobiaceae: Ostreobium | Ostreobium sp.R2 AR-2023 | Cnidaria:Scleractinea:Siderastreidae | *Siderastrea siderea* | 3/15/2020 | Colombia:Cartagena | 10.26N 75.61W | 10 |
| OQ935492 | ANDES-IM7781 | WP49 | Eukaryota:Viridiplantae:Chlorophyta: Ulvophyceae: Bryopsidales: Ostreobiaceae: Ostreobium | Ostreobium sp.E AR-2023 | Cnidaria:Scleractinea:Poritidae | *Porites sp.* | 5/12/2019 | Colombia:San Andres island | 12.51N 81.73W | 20 |
| OQ935493 | ANDES-IM7752 | WP16 | Eukaryota:Viridiplantae:Chlorophyta: Ulvophyceae: Bryopsidales: Ostreobiaceae: Ostreobium | Ostreobium sp.A AR-2023 | Cnidaria:Scleractinea:Merulinidae | *Orbicella annularis* | 5/12/2019 | Colombia:San Andres island | 12.51N 81.73W | 16 |
| OQ935494 | ANDES-IM7892 | CTG10 | Eukaryota:Viridiplantae:Chlorophyta: Ulvophyceae: Bryopsidales: Ostreobiaceae: Ostreobium | Ostreobium sp.F AR-2023 | Cnidaria:Scleractinea:Agariciidae | *Agaricia grahamae* | 3/15/2020 | Colombia:Cartagena | 10.26N 75.61W | 10 |
| OQ935495 | ANDES-IM7491 | I23 | Eukaryota:Viridiplantae:Chlorophyta: Ulvophyceae: Bryopsidales: Ostreobiaceae: Ostreobium | Ostreobium sp.E5 AR-2023 | Cnidaria:Scleractinea:Agariciidae | *Agaricia fragilis* | 6/14/2018 | Colombia:Cartagena | 10.26N 75.62W | 20 |
| OQ935496 | ANDES-IM7748 | WP12 | Eukaryota:Viridiplantae:Chlorophyta: Ulvophyceae: Bryopsidales: Ostreobiaceae: Ostreobium | Ostreobium sp.E3 AR-2023 | Cnidaria:Scleractinea:Agariciidae | *Helioseris cucullata* | 5/12/2019 | Colombia:San Andres island | 12.51N 81.73W | 15 |
| OQ935497 | ANDES-IM8205 | BT2 CTG48 | Eukaryota:Viridiplantae:Chlorophyta: Ulvophyceae: Bryopsidales: Ostreobiaceae: Ostreobium | Ostreobium sp.F AR-2023 | Cnidaria:Scleractinea:Agariciidae | *Agaricia tenuifolia* | 3/22/2022 | Colombia:Cartagena | 10.09N 75.87W | 15 |
| OQ935498 | ANDES-IM6417 | IMD78 | Eukaryota:Viridiplantae:Chlorophyta: Ulvophyceae: Bryopsidales: Ostreobiaceae: Ostreobium | Ostreobium sp.I AR-2023 | Cnidaria:Scleractinea:Agariciidae | *Agaricia lamarcki* | 10/9/2017 | Colombia:Cartagena | 10.27N 75.61W | 40 |
| OQ935499 | ANDES-IM7742 | WP06 | Eukaryota:Viridiplantae:Chlorophyta: Ulvophyceae: Bryopsidales: Ostreobiaceae: Ostreobium | Ostreobium sp.A AR-2023 | Cnidaria:Scleractinea:Agariciidae | *Agaricia agaricites* | 5/12/2019 | Colombia:San Andres island | 12.51N 81.73W | 15 |
| OQ935500 | ANDES-IM7510 | O3 | Eukaryota:Viridiplantae:Chlorophyta: Ulvophyceae: Bryopsidales: Ostreobiaceae: Ostreobium | Ostreobium sp.E5 AR-2023 | Cnidaria:Scleractinea:Merulinidae | *Orbicella faveolata* | 6/14/2018 | Colombia:Cartagena | 10.27N 75.61W | 25 |
| OQ935501 | ANDES-IM7867 | CTG20 | Eukaryota:Viridiplantae:Chlorophyta: Ulvophyceae: Bryopsidales: Ostreobiaceae: Ostreobium | Ostreobium sp.A2 AR-2023 | Cnidaria:Scleractinea:Poritidae | *Porites sp.* | 3/15/2020 | Colombia:Cartagena | 10.26N 75.61W | 10 |
| OQ935502 | ANDES-IM8209 | IT CTG53 | Eukaryota:Viridiplantae:Chlorophyta: Ulvophyceae: Bryopsidales: Ostreobiaceae: Ostreobium | Ostreobium sp.E4 AR-2023 | Cnidaria:Scleractinea:Agariciidae | *Agaricia lamarcki* | 3/22/2022 | Colombia:Cartagena | 10.14N 75.44W | 10 |
| OQ935503 | ANDES-IM7510 | T01 | Eukaryota:Viridiplantae:Chlorophyta: Ulvophyceae: Bryopsidales: Ostreobiaceae: Ostreobium | Ostreobium sp.R4 AR-2023 | Cnidaria:Scleractinea:Agariciidae | *Agaricia lamarcki* | 6/14/2018 | Colombia:Cartagena | 10.27N 75.63W | 25 |
| OQ935504 | ANDES-IM7868 | CTG09 | Eukaryota:Viridiplantae:Chlorophyta: Ulvophyceae: Bryopsidales: Ostreobiaceae: Ostreobium | Ostreobium sp.E5 AR-2023 | Cnidaria:Scleractinea:Pocilloporidae | *Madracis sp.* | 3/15/2020 | Colombia:Cartagena | 10.26N 75.61W | 10 |
| OQ935505 | ANDES-IM7754 | WP18 | Eukaryota:Viridiplantae:Chlorophyta: Ulvophyceae: Bryopsidales: Ostreobiaceae: Ostreobium | Ostreobium sp.E3 AR-2023 | Cnidaria:Scleractinea:Agariciidae | *Agaricia humilis* | 5/12/2019 | Colombia:San Andres island | 12.51N 81.73W | 8 |
| OQ935506 | ANDES-IM8202 | BT2 CTG32 | Eukaryota:Viridiplantae:Chlorophyta: Ulvophyceae: Bryopsidales: Ostreobiaceae: Ostreobium | Ostreobium sp.F AR-2023 | Cnidaria:Scleractinea:Agariciidae | *Agaricia tenuifolia* | 3/22/2022 | Colombia:Cartagena | 10.09N 75.87W | 15 |
| OQ935507 | ANDES-IM7860 | CTG48 | Eukaryota:Viridiplantae:Chlorophyta: Ulvophyceae: Bryopsidales: Ostreobiaceae: Ostreobium | Ostreobium sp.A AR-2023 | Cnidaria:Scleractinea:Agariciidae | *Helioseris cucullata* | 3/15/2020 | Colombia:Cartagena | 10.26N 75.62W | 33 |
| OQ935508 | ANDES-IM8200 | BT CTG30 | Eukaryota:Viridiplantae:Chlorophyta: Ulvophyceae: Bryopsidales: Ostreobiaceae: Ostreobium | Ostreobium sp.E5 AR-2023 | Cnidaria:Scleractinea:Agariciidae | *Agaricia fragilis* | 3/22/2022 | Colombia:Cartagena | 10.09N 75.87W | 10 |
| OQ935509 | ANDES-IM8203 | BT2 CTG41 | Eukaryota:Viridiplantae:Chlorophyta: Ulvophyceae: Bryopsidales: Ostreobiaceae: Ostreobium | Ostreobium sp.E5 AR-2023 | Cnidaria:Scleractinea:Agariciidae | *Agaricia grahamae* | 3/22/2022 | Colombia:Cartagena | 10.09N 75.87W | 15 |
| OQ935510 | ANDES-IM7757 | WP22 | Eukaryota:Viridiplantae:Chlorophyta: Ulvophyceae: Bryopsidales: Ostreobiaceae: Ostreobium | Ostreobium sp.E3 AR-2023 | Cnidaria:Scleractinea:Astrocoeniidae | *Stephanocoenia intersepta* | 5/12/2019 | Colombia:San Andres island | 12.51N 81.73W | 10 |
| OQ935511 | ANDES-IM7891 | CTG18 | Eukaryota:Viridiplantae:Chlorophyta: Ulvophyceae: Bryopsidales: Ostreobiaceae: Ostreobium | Ostreobium sp.E2 AR-2023 | Cnidaria:Scleractinea:Siderastreidae | *Siderastrea siderea* | 3/15/2020 | Colombia:Cartagena | 10.26N 75.61W | 10 |
| OQ935512 | ANDES-IM7890 | CTG04 | Eukaryota:Viridiplantae:Chlorophyta: Ulvophyceae: Bryopsidales: Ostreobiaceae: Ostreobium | Ostreobium sp.R4 AR-2023 | Cnidaria:Scleractinea:Agariciidae | *Agaricia grahamae* | 3/15/2020 | Colombia:Cartagena | 10.26N 75.61W | 10 |
| OQ935513 | ANDES-IM8211 | SH 02 | Eukaryota:Viridiplantae:Chlorophyta: Ulvophyceae: Bryopsidales: Ostreobiaceae: Ostreobium | Ostreobium sp.E3 AR-2023 | Cnidaria:Scleractinea:Montastraeidae | *Montrastea cavernosa* | 3/22/2022 | Colombia:San Andres island | 12.51N 81.73W | 7 |
| OQ935514 | ANDES-IM6528 | OCD01 | Eukaryota:Viridiplantae:Chlorophyta: Ulvophyceae: Bryopsidales: Ostreobiaceae: Ostreobium | Ostreobium sp.B3 AR-2023 | Cnidaria:Scleractinea:Agariciidae | *Agaricia lamarcki* | 9/10/2017 | Colombia:Cartagena | 10.27N 75.61W | 33 |
| OQ935515 | ANDES-IM7480 | I10 | Eukaryota:Viridiplantae:Chlorophyta: Ulvophyceae: Bryopsidales: Ostreobiaceae: Ostreobium | Ostreobium sp.E AR-2023 | Cnidaria:Scleractinea:Agariciidae | *Agaricia humilis* | 6/14/2018 | Colombia:Cartagena | 10.26N 75.62W | 20 |
| OQ935516 | ANDES-IM7512 | O7 | Eukaryota:Viridiplantae:Chlorophyta: Ulvophyceae: Bryopsidales: Ostreobiaceae: Ostreobium | Ostreobium sp.E AR-2023 | Cnidaria:Scleractinea:Merulinidae | *Orbicella faveolata* | 6/14/2018 | Colombia:Cartagena | 10.27N 75.61W | 25 |
| OQ935517 | ANDES-IM7499 | M7 | Eukaryota:Viridiplantae:Chlorophyta: Ulvophyceae: Bryopsidales: Ostreobiaceae: Ostreobium | Ostreobium sp.E AR-2023 | Cnidaria:Scleractinea:Merulinidae | *Orbicella annularis* | 6/14/2018 | Colombia:Cartagena | 10.26N 75.62W | 25 |
| OQ935518 | ANDES-IM7759 | WP24 | Eukaryota:Viridiplantae:Chlorophyta: Ulvophyceae: Bryopsidales: Ostreobiaceae: Ostreobium | Ostreobium sp.E3 AR-2023 | Cnidaria:Scleractinea:Agariciidae | *Agaricia agaricites* | 5/12/2019 | Colombia:San Andres island | 12.51N 81.73W | 10 |
| OQ935519 | ANDES-IM7896 | CTG11 | Eukaryota:Viridiplantae:Chlorophyta: Ulvophyceae: Bryopsidales: Ostreobiaceae: Ostreobium | Ostreobium sp.A AR-2023 | Cnidaria:Scleractinea:Agariciidae | *Helioseris cucullata* | 3/15/2020 | Colombia:Cartagena | 10.26N 75.61W | 10 |
| OQ935520 | ANDES-IM7871 | CTG25 | Eukaryota:Viridiplantae:Chlorophyta: Ulvophyceae: Bryopsidales: Ostreobiaceae: Ostreobium | Ostreobium sp.E5 AR-2023 | Cnidaria:Scleractinea:Agariciidae | *Helioseris cucullata* | 3/15/2020 | Colombia:Cartagena | 10.26N 75.61W | 10 |
| OQ935521 | ANDES-IM7511 | O5 | Eukaryota:Viridiplantae:Chlorophyta: Ulvophyceae: Bryopsidales: Ostreobiaceae: Ostreobium | Ostreobium sp.E AR-2023 | Cnidaria:Scleractinea:Merulinidae | *Orbicella faveolata* | 6/14/2018 | Colombia:Cartagena | 10.27N 75.61W | 25 |
| OQ935522 | ANDES-IM7777 | WP45 | Eukaryota:Viridiplantae:Chlorophyta: Ulvophyceae: Bryopsidales: Ostreobiaceae: Ostreobium | Ostreobium sp.E3 AR-2023 | Cnidaria:Scleractinea:Agariciidae | *Agaricia grahamae* | 5/12/2019 | Colombia:San Andres island | 12.51N 81.73W | 17 |
| OQ935523 | ANDES-IM8204 | BT2 CTG44 | Eukaryota:Viridiplantae:Chlorophyta: Ulvophyceae: Bryopsidales: Ostreobiaceae: Ostreobium | Ostreobium sp.F AR-2023 | Cnidaria:Scleractinea:Agariciidae | *Agaricia humilis* | 3/22/2022 | Colombia:Cartagena | 10.09N 75.87W | 15 |
| OQ935524 | ANDES-IM7495 | M3 | Eukaryota:Viridiplantae:Chlorophyta: Ulvophyceae: Bryopsidales: Ostreobiaceae: Ostreobium | Ostreobium sp.E1 AR-2023 | Cnidaria:Scleractinea:Merulinidae | *Orbicella faveolata* | 6/14/2018 | Colombia:Cartagena | 10.26N 75.62W | 25 |
| OQ935525 | ANDES-IM8206 | IT CTG50 | Eukaryota:Viridiplantae:Chlorophyta: Ulvophyceae: Bryopsidales: Ostreobiaceae: Ostreobium | Ostreobium sp.E4 AR-2023 | Cnidaria:Scleractinea:Agariciidae | *Agaricia fragilis* | 3/22/2022 | Colombia:Cartagena | 10.14N 75.44W | 10 |
| OQ935526 | ANDES-IM7893 | CTG12 | Eukaryota:Viridiplantae:Chlorophyta: Ulvophyceae: Bryopsidales: Ostreobiaceae: Ostreobium | Ostreobium sp.F AR-2023 | Cnidaria:Scleractinea:Agariciidae | *Agaricia tenuifolia* | 3/15/2020 | Colombia:Cartagena | 10.26N 75.61W | 10 |
| OQ935527 | ANDES-IM7494 | M2 | Eukaryota:Viridiplantae:Chlorophyta: Ulvophyceae: Bryopsidales: Ostreobiaceae: Ostreobium | Ostreobium sp.E AR-2023 | Cnidaria:Scleractinea:Merulinidae | *Orbicella faveolata* | 6/14/2018 | Colombia:Cartagena | 10.26N 75.62W | 25 |
| OQ935528 | ANDES-IM7877 | CTG00 | Eukaryota:Viridiplantae:Chlorophyta: Ulvophyceae: Bryopsidales: Ostreobiaceae: Ostreobium | Ostreobium sp.E5 AR-2023 | Cnidaria:Scleractinea:Merulinidae | *Orbicella sp.* | 3/15/2020 | Colombia:Cartagena | 10.26N 75.61W | 10 |
| OQ935529 | ANDES-IM6394 | IMD2 | Eukaryota:Viridiplantae:Chlorophyta: Ulvophyceae: Bryopsidales: Ostreobiaceae: Ostreobium | Ostreobium sp.L1 AR-2023 | Cnidaria:Scleractinea:Agariciidae | *Agaricia lamarcki* | 10/9/2017 | Colombia:Cartagena | 10.27N 75.61W | 40 |
| OQ935530 | ANDES-IM8210 | IT CTG54 | Eukaryota:Viridiplantae:Chlorophyta: Ulvophyceae: Bryopsidales: Ostreobiaceae: Ostreobium | Ostreobium sp.E4 AR-2023 | Cnidaria:Scleractinea:Agariciidae | *Agaricia lamarcki* | 3/22/2022 | Colombia:Cartagena | 10.14N 75.44W | 10 |
| OQ935531 | ANDES-IM7760 | WP25 | Eukaryota:Viridiplantae:Chlorophyta: Ulvophyceae: Bryopsidales: Ostreobiaceae: Ostreobium | Ostreobium sp.E3 AR-2023 | Cnidaria:Scleractinea:Astrocoeniidae | *Stephanocoenia intersepta* | 5/12/2019 | Colombia:San Andres island | 12.51N 81.73W | 8 |
| OQ935532 | ANDES-IM7739 | WP01 | Eukaryota:Viridiplantae:Chlorophyta: Ulvophyceae: Bryopsidales: Ostreobiaceae: Ostreobium | Ostreobium sp.F AR-2023 | Cnidaria:Scleractinea:Agariciidae | *Agaricia agaricites* | 5/12/2019 | Colombia:San Andres island | 12.51N 81.73W | 15 |
| OQ935533 | ANDES-IM8208 | IT CTG52 | Eukaryota:Viridiplantae:Chlorophyta: Ulvophyceae: Bryopsidales: Ostreobiaceae: Ostreobium | Ostreobium sp.E4 AR-2023 | Cnidaria:Scleractinea:Agariciidae | *Agaricia agaricites* | 3/22/2022 | Colombia:Cartagena | 10.14N 75.44W | 10 |
| OQ935534 | ANDES-IM7493 | M1 | Eukaryota:Viridiplantae:Chlorophyta: Ulvophyceae: Bryopsidales: Ostreobiaceae: Ostreobium | Ostreobium sp.E AR-2023 | Cnidaria:Scleractinea:Agariciidae | *Agaricia agaricites* | 6/14/2018 | Colombia:Cartagena | 10.26N 75.62W | 25 |
|  | ANDES-IM7950 | I6 | Eukaryota:Viridiplantae:Chlorophyta: Ulvophyceae: Bryopsidales: Ostreobiaceae: Ostreobium | Ostreobium sp. F | Cnidaria:Scleractinea:Poritidae | *Porites astreoides* | 11/26/2020 | Colombia:Cartagena | 10.26N 75.62W | 20 |
|  | ANDES-IM8041 | J2 | Eukaryota:Viridiplantae:Chlorophyta: Ulvophyceae: Bryopsidales: Ostreobiaceae: Ostreobium | Ostreobium sp. F | Cnidaria:Scleractinea:Poritidae | *Porites porites* | 6/26/2019 | Colombia:San Andres island | 12.51N 81.73W | 20 |
|  | ANDES-IM8013 | 44 | Eukaryota:Viridiplantae:Chlorophyta: Ulvophyceae: Bryopsidales: Ostreobiaceae: Ostreobium | Ostreobium sp. G | Cnidaria:Scleractinea:Poritidae | *Porites furcata* | 11/26/2020 | Colombia:Cartagena | 10.26N 75.62W | 33 |
|  | ANDES-IM7944 | I10 | Eukaryota:Viridiplantae:Chlorophyta: Ulvophyceae: Bryopsidales: Ostreobiaceae: Ostreobium | Ostreobium sp. P4c | Cnidaria:Scleractinea:Poritidae | *Porites colonensis* | 11/26/2020 | Colombia:Cartagena | 10.26N 75.62W | 20 |
|  | ANDES-IM8049 | M4 | Eukaryota:Viridiplantae:Chlorophyta: Ulvophyceae: Bryopsidales: Ostreobiaceae: Ostreobium | Ostreobium sp. P6c | Cnidaria:Scleractinea:Poritidae | *Porites colonensis* | 6/26/2019 | Colombia:San Andres island | 12.51N 81.73W | 8 |
|  | ANDES-IM8043 | L3 | Eukaryota:Viridiplantae:Chlorophyta: Ulvophyceae: Bryopsidales: Ostreobiaceae: Ostreobium | Ostreobium sp. Q | Cnidaria:Scleractinea:Poritidae | *Porites furcata* | 6/26/2019 | Colombia:San Andres island | 12.51N 81.73W | 35 |
|  | ANDES-IM8006 | 22 | Eukaryota:Viridiplantae:Chlorophyta: Ulvophyceae: Bryopsidales: Ostreobiaceae: Ostreobium | Ostreobium sp. A2 | Cnidaria:Scleractinea:Poritidae | *Porites astreoides* | 11/26/2020 | Colombia:Cartagena | 10.27N 75.61W | 18 |

**Supplementary Table 2.** Complementary rbcL sequences accession numbers used in the present study.

| **Author** | **NCBI Code** | **Ostreobium clade** |
| --- | --- | --- |
| Gonzalez-Zapata et al. 2018 | MF135637.1 | Ostreobium sp. D1 |
| Gonzalez-Zapata et al. 2018 | MF135632.1 | Ostreobium sp. D3 |
| Gonzalez-Zapata et al. 2018 | MF135629.1 | Ostreobium sp. E |
| Gonzalez-Zapata et al. 2018 | MF135635.1 | Ostreobium sp. H2 |
| Gonzalez-Zapata et al. 2018 | MF135639.1 | Ostreobium sp. I |
| Gonzalez-Zapata et al. 2018 | MF135636.1 | Ostreobium sp. J |
| Gonzalez-Zapata et al. 2018 | MF135638.1 | Ostreobium sp. K |
| Gonzalez-Zapata et al. 2018 | MF135640.1 | Ostreobium sp. L |
| Gonzalez-Zapata et al. 2018 | MF135643.1 | Ostreobium sp. N |
| Gonzalez-Zapata et al. 2018 | MF135628.1 | Ostreobium sp. O |
| Gonzalez-Zapata et al. 2018 | MF135641.1 | Ostreobium sp. M |
| Gonzalez-Zapata et al. 2018 | MF135631.1 | Ostreobium sp. D2 |
| Gonzalez-Zapata et al. 2018 | MF135634.1 | Ostreobium sp. H12 |
| Gonzalez-Zapata et al. 2018 | MF135627.1 | Ostreobium sp. I |
| Gonzalez-Zapata et al. 2018 | MF135626.1 | Ostreobium sp. I |
| Gonzalez-Zapata et al. 2018 | MF135625.1 | Ostreobium sp. H3 |
| Gonzalez-Zapata et al. 2018 | MF135624.1 | Ostreobium sp. H3 |
| Gonzalez-Zapata et al. 2018 | MF135623.1 | Ostreobium sp. N |
| Gonzalez-Zapata et al. 2018 | MF135622.1 | Ostreobium sp. I |
| Direct submission O'Kelly, C.J., Mottet, G.J., Santoni, S. and Tribollet, A. 2011 | KC685549.1 | Ostreobium sp. TeF |
| Masse et al. 2018 | MG570002 | Ostreobium sp. K |
| Masse et al. 2018 | MG570016 | Ostreobium sp. P4 |
| Masse et al. 2018 | MG569998 | Ostreobium sp. P3 |
| Masse et al. 2018 | MG570015 | Ostreobium sp. P4 |
| Masse et al. 2018 | MG569992.1 | Ostreobium sp. P5 |
| Masse et al. 2018 | MG569989 | Ostreobium sp. P6 |
| Masse et al. 2018 | MK095219.1 | Ostreobium sp. P1 |
| Gutner-Hoch & Fine 2011 | JF801728.1 | Ostreobium queketii |
| Gutner-Hoch & Fine 2011 | JF801728.1 | Ostreobium clade A |
| Gutner-Hoch & Fine 2011 | JF801729.1 | Ostreobium clade B |
| Gutner-Hoch & Fine 2011 | JF801730.1 | Ostreobium clade C |
| Gutner-Hoch & Fine 2011 | JF801731.1 | Ostreobium clade D |
| Gutner-Hoch & Fine 2011 | JF801732.1 | Ostreobium clade E |
| Gutner-Hoch & Fine 2011 | JF801733.1 | Ostreobium clade F |
| Gutner-Hoch & Fine 2011 | JF801734.1 | Ostreobium clade G |
| Direct submission Cohen,E., Tikochinski,Y. and Fine,M. | KT279998 | Ostreobium sp. A |
| Direct submission Cohen,E., Tikochinski,Y. and Fine,M. | KT280003 | Ostreobium sp. F |
| Direct submission Cohen,E., Tikochinski,Y. and Fine,M. | KT795105 | Ostreobium sp. J |
| Direct submission | MG753774.1 | Caulerpa lentillifera f. tomentella |
| Direct submission | AB054017.1 | Caulerpa webbiana |
| Direct submission Cohen,E., Tikochinski,Y. and Fine,M. | KT280002.1 | Ostreobium sp. E |
| Direct submission Cohen,E., Tikochinski,Y. and Fine,M. | KT280004 | Ostreobium sp. I |
| Direct submission Cohen,E., Tikochinski,Y. and Fine,M. | KT795106.1 | Ostreobium sp. B2 |
| Direct submission O'Kelly, C.J., Mottet, G.J., Santoni, S. and Tribollet, A. 2011 | KC685544.1 | Ostreobium sp.TeG strain FHL094 |

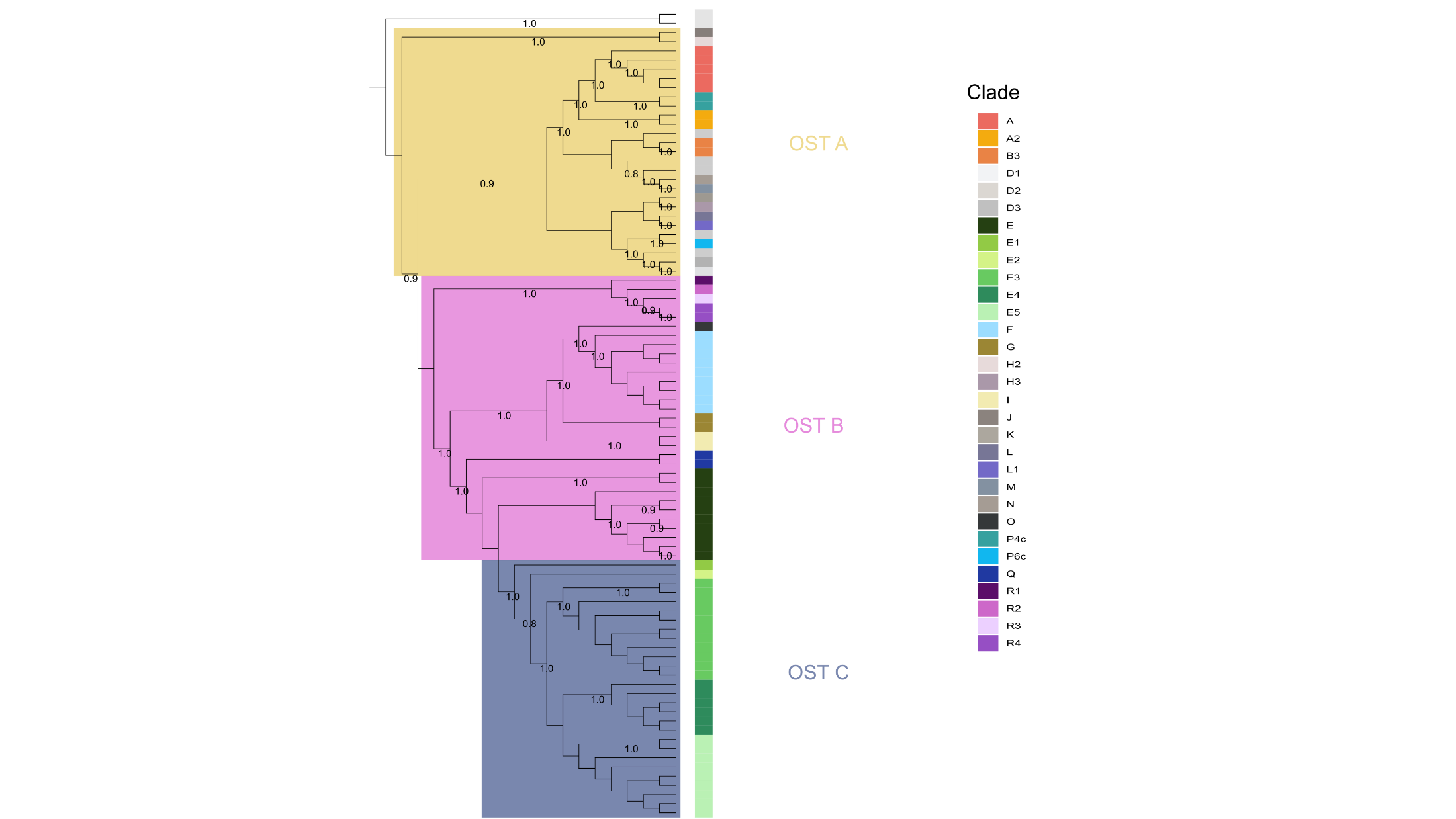
**Supplementary Figure A**. Phylogenetic tree indicating the three ‘superclades’ of *Ostreobium.*
